# What microbes want: exploring microbial substrate preferences with the Web of Microbes Agent

**DOI:** 10.64898/2026.02.23.707520

**Authors:** Trent R. Northen, Markus de Raad, Suzanne M. Kosina, Peter F. Andeer, Vlastimil Novak, Bradley Biggs, Hao Peng, Timothy Paulitz, Adam P. Arkin, Katherine B. Louie, Mingxun Wang, Benjamin P. Bowen

## Abstract

Understanding and predicting bacterial substrate preferences has broad utility from microbial interactions to selecting prebiotics. Isolate exometabolite profiling directly measures which compounds a given microbe utilizes from an array of metabolites in the environment. However, modeling, mining, and integrating these data are challenging. Here, we introduce a Bayesian Personalized Ranking (BPR) model applied to substrate preferences which we find learns to rank compounds by a given microbe’s preference. It was found to outperform the other ranking models (AUC = 0.93), proved robust to ablation, showed strong within-genus isolate pairs correlation (Spearman rank = 0.78) and predictive ability for new data. BPR was then used to create the Web of Microbes (WoM) Agent by integrating it with the Phydon growth model and Large Language Model (LLM) for autonomous orchestration tool calling and analysis. The WoM Agent accurately predicted substrate consumption by existing strain grown on a novel medium and correctly identified bacteria enriched in soil metabolite spike-in experiments. Additionally, the WoM Agent can use autonomous reasoning including to predict substrates that will selectively promote the growth of one clade of bacteria over another including helping interpret results and suggest new hypotheses and experiments. We anticipate broad applications in microbial cultivation, microbiome engineering, and environmental microbiology, with the agent’s capabilities further extensible through the integration of additional tools and use of rapidly improving LLMs.

## Introduction

Tools for predicting and understanding microbial substrate preferences can inform interventions in important applications, including low-input agriculture, bioproducts, human and animal health, and environmental management. A large body of work has demonstrated the value of genome-scale modeling, consumer-resource models, hybrid models, and trait-based models for predicting microbial substrate preferences, among other approaches^1–4^. However, these models generally require substantial additional data beyond direct measurements of substrate use, such annotated genomes, biomass measurements, genome based trait predictions, etc.^1,3^. Furthermore, modeling, interpreting, and integrating substrate preference are currently laborious processes and generally performed by subject experts^2,5^.

The integration of Large Language Models (LLMs) into scientific workflows has catalyzed the development of specialized agents capable of reasoning, planning, and tool execution across chemical and biological domains including expanding access to state-of-the-art models. Agents like ChemCrow have augmented LLMs with expert tools to plan and execute chemical synthesis and material design experiments^6,7^. Talk2Biomodels (T2B) has democratized access to systems biology by enabling natural language interaction with kinetic ordinary differential equation (ODE) models ^8^. Simulation-augmented graph neural networks like SIMBA have integrated metabolomics and metabolic network simulations to predict microbial growth and community composition ^9^. These advances suggest that pairing domain-specific models with LLM-driven interfaces could make complex microbial analyses more accessible and interpretive, yet no existing agent addresses substrate preference modeling.

Substrate preferences can be measured experimentally using mass spectrometry to quantify the selective depletion of metabolites by microbes from complex media ^10^. This exometabolite profiling approach has proven useful across microbial genomics, ecology, and interaction studies. For example, Kell and co-workers showed that metabolite use profiles alone provided mechanistic insights into yeast mutant phenotypes ^10^. Exometabolite profiling can also provide ecology insights, for example application to soil biocrust microbes revealed largely non-overlapping substrate preferences, providing insights into community assembly ^11,12^. The approach has also been applied to pathogen diagnostics and microbiome engineering ^13–15^.

Despite the utility of exometabolite profiling, existing approaches for modeling substrate preferences — including genome-scale modeling^5^, Lotka-Volterra models^16^, and consumer-resource models^2^, etc — are active areas of research and typically require additional data beyond the metabolite profiles themselves. Since exometabolite profiling fundamentally measures substrate preferences, we hypothesized that ranking algorithms developed for recommender systems (e.g. online shopping), which learn to rank items by human preference, could be naturally adapted to rank substrates by microbial preference^17,18^. For example, where a recommender system learns that a user prefers certain videos over others from implicit feedback, a ranking model can learn that a microbe preferentially depletes certain substrates over others from exometabolite profiles. The resulting preference model could be run using a LLM to make the model accessible to a broad range of users and use cases.

Here, we describe the Web of Microbes Agent (WoM Agent), currently available at http://webofmicrobes.org/, which combines an LLM with a Bayesian Personalized Ranking (BPR) model of substrate preferences and a growth model. This ranking model was selected after comparing the ranking algorithms described by Rendle et al.^17^ on their ability to accurately predict taxonomic patterns in substrate preferences. BPR was found to outperform the other approaches, proved robust to ablation, was validated using new data, and was therefore selected for integration with the LLM along with the open-source Phydon growth model^19^. We tested WoM’s ability to predict the substrate preferences of new isolates and media; evaluated its responses against published results; evaluated whether the agent could predict which bacteria would be promoted by the addition of metabolites spiked-in to soil; and evaluated WoM’s capacity for autonomous reasoning on complex interpretive tasks.

## Results

### Bayesian Personalized Ranking effectively learns bacteria substrate preferences

Our overall objective was to develop an agent that leverages WoM-formatted exometabolite data for predicting and exploring microbial substrate preferences (**Fig. 1**). First we needed to develop a substrate preferences model. To train this model, we used data and the data format from our previous publication^15^ including a total of 226 isolates and 119 metabolites resulting from exometabolite profiling experiments comparing inoculated vs. non-inoculated media. Briefly, to format the data significantly altered metabolites were assigned a score from −5 (strongly consumed) to 5 (strongly produced) based on log2 fold change as previously described ^15^.

**Figure 1.**
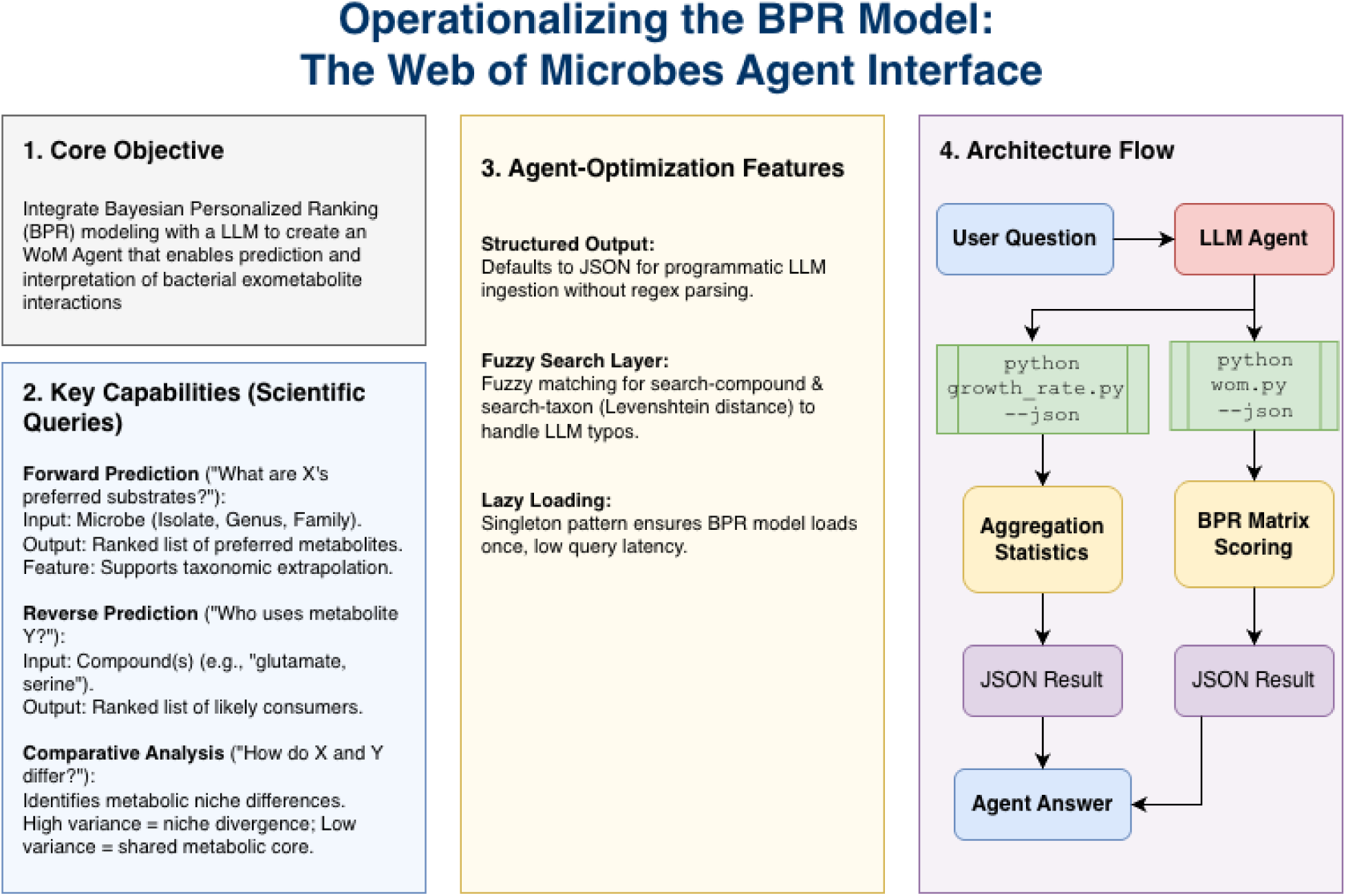
Operational architecture of the Web of Microbes Agent. The system integrates a Bayesian Personalized Ranking (BPR) consumption model with a Large Language Model (LLM) agent. This enables bidirectional predictive queries (microbe-to-metabolite and metabolite-to-microbe) and comparative metabolic-niche analysis through an architecture optimized for robust, low latency interaction.

We next used these data to evaluate the potential of three ranking-based algorithms for learning bacterial substrate preferences from these WoM-formatted data (**Fig. 2**). The first two are factorization methods: truncated Singular Value Decomposition (SVD) and Weighted Regularized Matrix Factorization (WR-MF)^20,21^. The third is a pairwise optimization algorithm, Bayesian Personalized Ranking (BPR)^17^. Briefly, truncated SVD uses linear algebra to reduce noise and uncover latent patterns in explicit preference scores. WR-MF extends this to implicit data by assigning varying confidence levels to observed versus missing interactions. In contrast, BPR optimizes for the relative preference between pairs of substrates, prioritizing compounds a microbe depletes over those it leaves unaffected.

**Figure 2.**
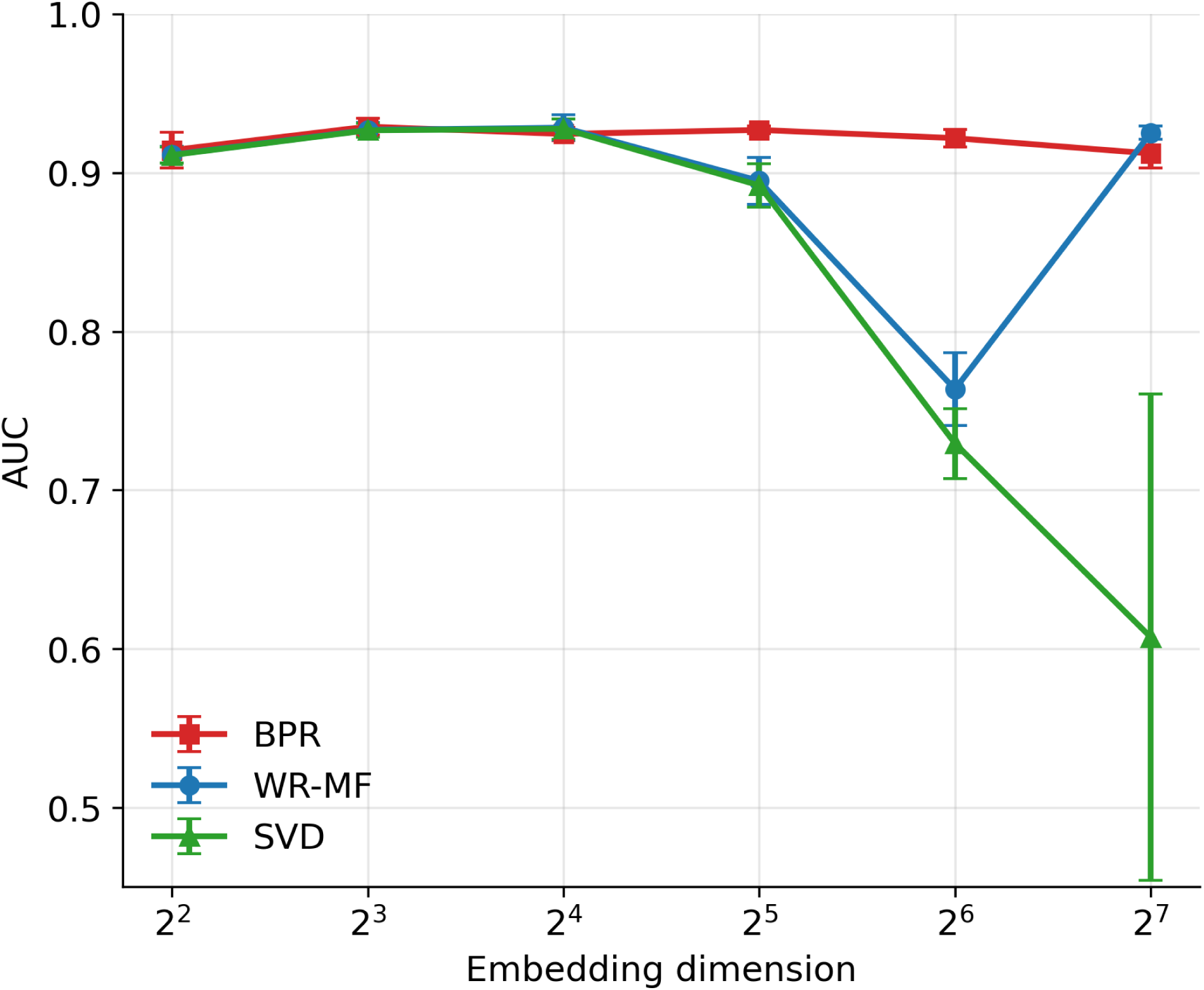
Benchmark evaluation of metabolite consumption prediction. Leave-one-out cross-validation comparing Bayesian Personalized Ranking (BPR), Weighted Regularized Matrix Factorization (WR-MF), and truncated Singular Value Decomposition (SVD) across embedding dimensions from 4 to 128. Area under the ROC curve (AUC) showing the ability to rank held-out positive metabolites above unobserved metabolites where 0.5 is random ranking.

To select the final model for integration with a LLM agent we compared the performance of these models using leave-one-out cross-validation to assess model robustness. As shown in **Fig. 2**, BPR maintained stable performance across all embedding dimensions (AUC ≈ 0.93), while SVD and WR-MF degraded substantially at higher dimensions, likely due to overfitting. All three methods performed equivalently at low dimensions (4–16), suggesting the underlying consumption signal is low-rank. Based on this comparison, we selected BPR for all subsequent analyses. We used 32-dimensional embeddings for all subsequent analyses.

### Substrate preferences are generally conserved at the genus level

To examine the extent that substrate preferences and phylogenetically conserved we compared substrate profiles of microbes at different taxonomic levels. **Fig. 3** examines the distances of microbial substrate profiles at the genus, family, order and class levels. Visual examination of the histograms comparing intra- and inter-genus similarities in substrate preference profiles clearly show that microbes of the same genus as a whole have more similar profiles than microbes from other genera. This trend fades at broader taxonomic comparisons so that at the class level the two distributions almost overlap. Isolates belonging to the same genus had substantially higher cosine similarity in the BPR embedding space than isolates from different genera (within-genus: 0.53 ± 0.24; between-genus: 0.23 ± 0.22; Mann-Whitney U test, *p* < 10⁻³⁰⁰). Genus identity explained 42% of the total variance in embedding space (PERMANOVA pseudo-F = 3.21, p = 0.001, 999 permutations). This conservation was also evident in metabolite preference rankings: within-genus isolate pairs showed a mean Spearman rank correlation of 0.78 ± 0.24 compared to 0.56 ± 0.29 for between-genus pairs (Mann-Whitney *p* < 10⁻¹⁹⁸). These correlations included 188 isolates across 35 genera with at least two representatives each.

**Figure 3.**
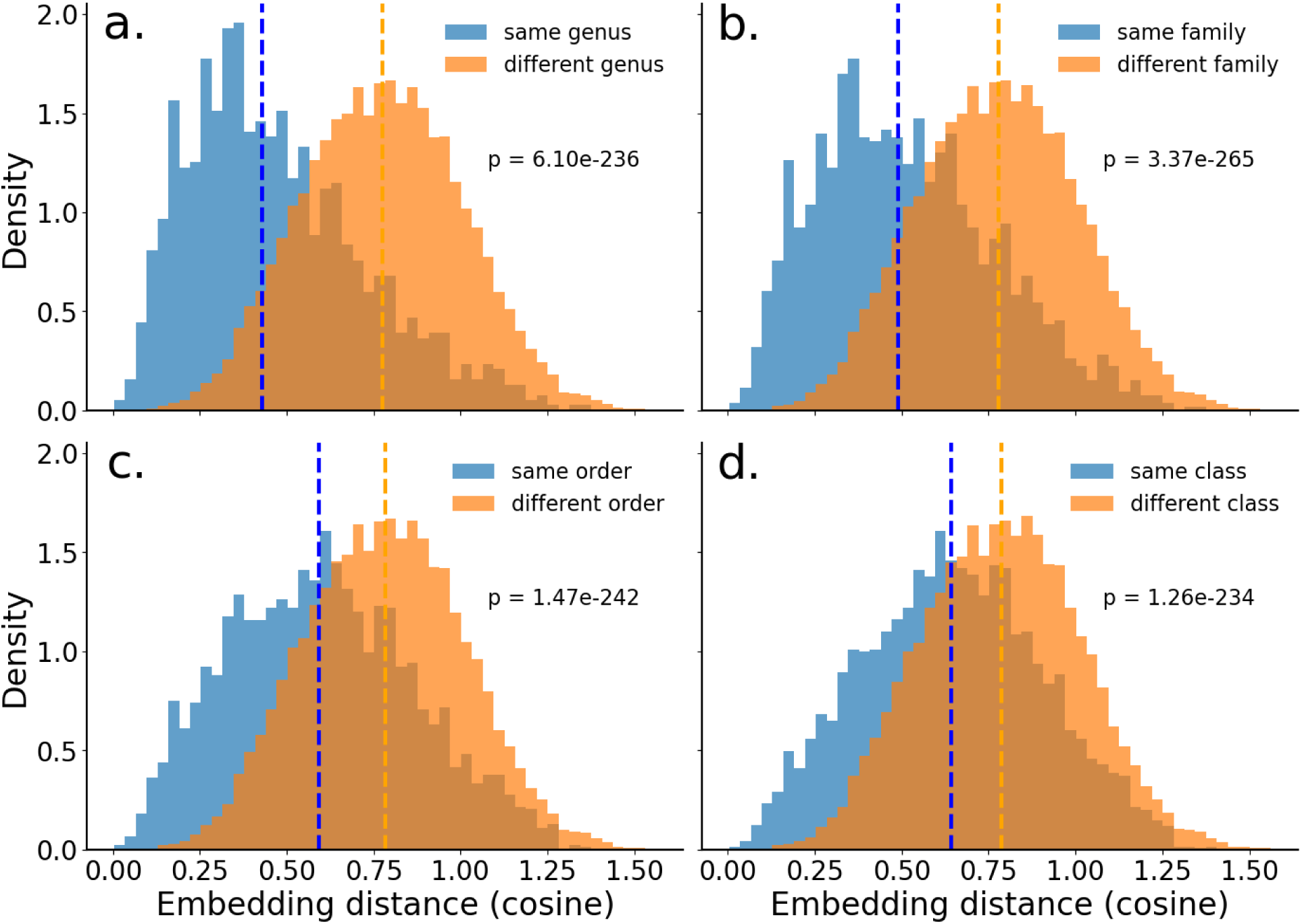
Substrate profiles are more similar at the genus level and less distinct at higher taxonomic levels. BPR scores are calculated from the dot product of the bacterial embedding vector with each metabolite embedding vector. These embedding vectors can be used to efficiently calculate distance between microbes of the same (a) genus, (b) family, (c) order, and (d) class. This plot clearly shows the agreement with embedding distance and phylogeny especially for closely related isolates (dashed-line: median value; independent t-tests < 0.05).

We next examined if the model accurately captured experimental substrate preferences (**Fig. 4**). Globally, metabolite preferences (i.e., BPR score) across all of the isolates show a clear left-to-right transition from blue (consumed) to pink (neutral) across nearly all isolates. This indicates that the BPR model is successfully ranking consumed metabolites higher than non-consumed ones and the model occasionally exhibits inferred preference. Notably, the model occasionally ranks experimentally identified neutral metabolites above consumed ones, an implicit ranking behavior intrinsic to BPR’s pairwise optimization. In other words, this is a *de novo* prediction of substrate preferences never directly measured. This emergent property transforms BPR from a summarizer of known consumption patterns into a hypothesis generator, pointing to specific untested microbe–metabolite interactions that could be rapidly validated to expand exometabolite reference datasets. To identify metabolites whose consumption distinguishes specific taxa, we computed distinctive scores for each genus–metabolite pair by projecting genus-level mean embeddings orthogonally to the global consumption direction. Metabolites consumed by more than 70% of all isolates were excluded due to being universally preferred, and the 60 remaining metabolites with the highest variance in distinctive score across genera were retained for visualization (**Fig. 5**).

**Figure 4.**
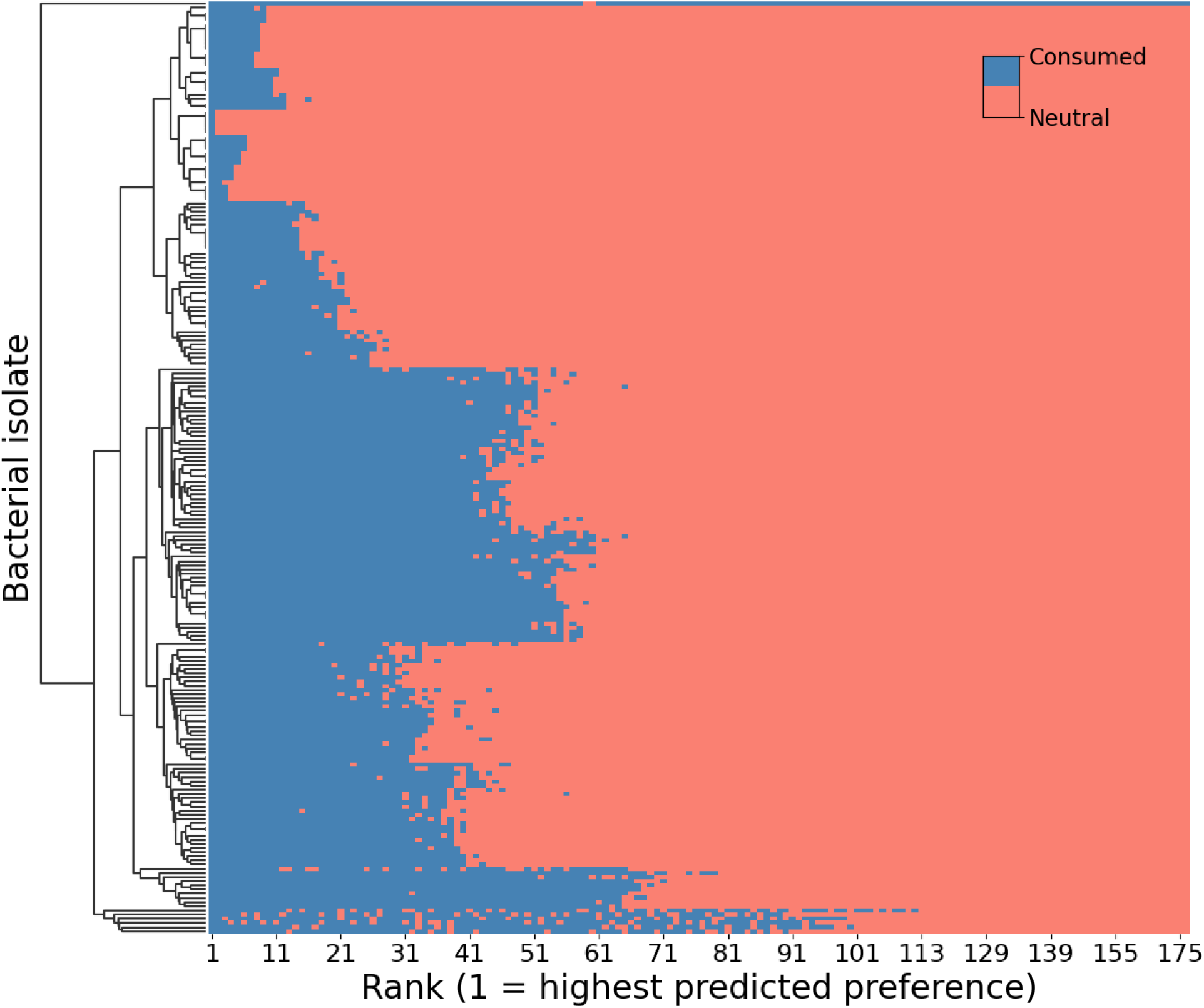
BPR model predictions concentrate consumed metabolites at top-ranked positions. Each row represents a bacterial isolate and each column a rank position, where rank 1 corresponds to the metabolite with the highest model-predicted preference score. Blue cells indicate experimentally confirmed consumption; salmon cells indicate no observed consumption. Rows are hierarchically clustered (Ward’s method) by their consumption-rank profiles, revealing groups of isolates with similar metabolic breadth. The strong concentration of blue toward the left side of the figure demonstrates that the model reliably assigns high ranks to consumed metabolites and the occasional interleaved salmon pixels illustrate the implicit ranking capabilities of the BPR model.

**Figure 5.**
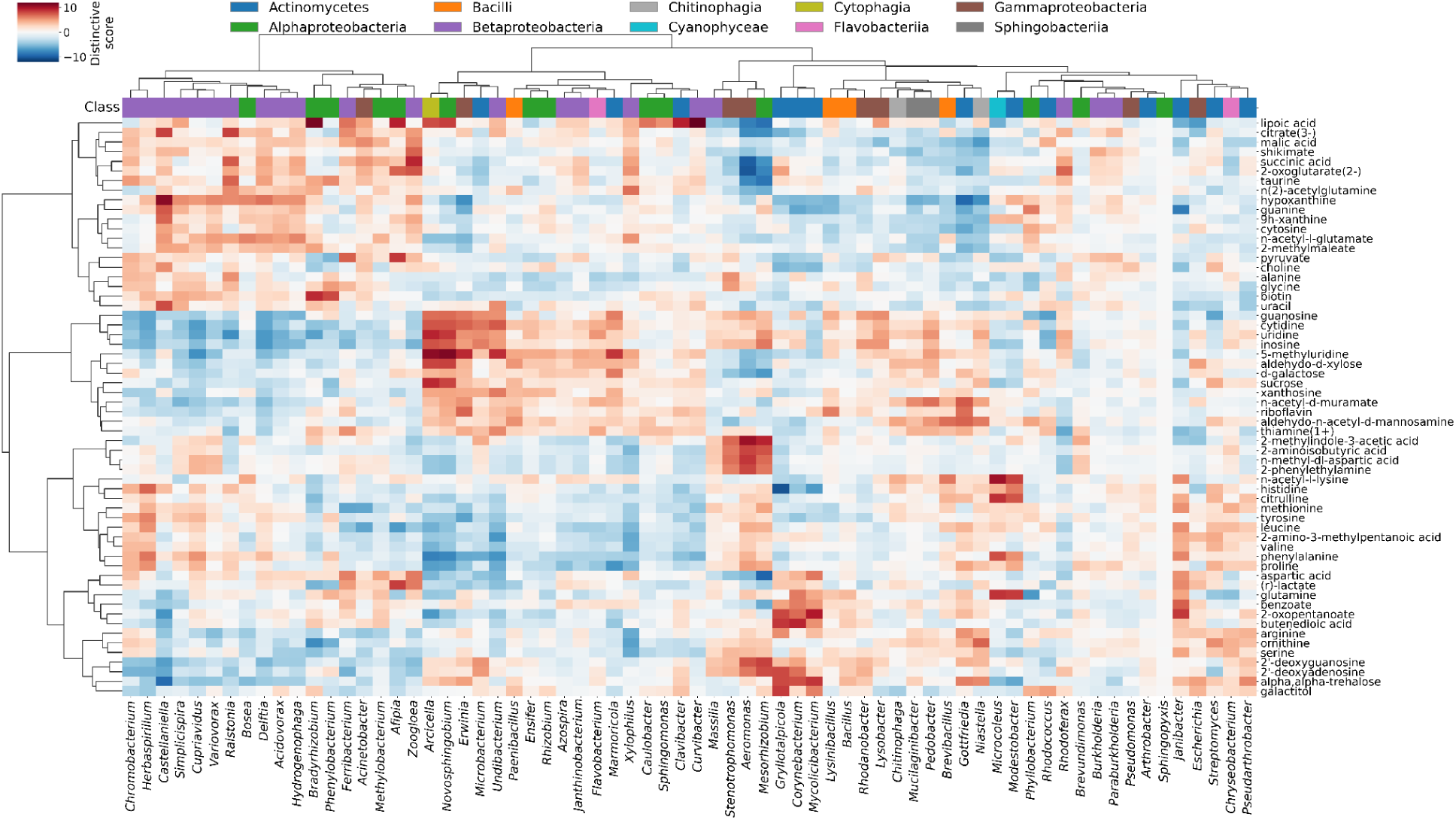
Genus-level distinctive metabolite preference heatmap. Distinctive scores for each genus-metabolite pair, showing the 60 metabolites with the highest distinctive score across genera. Genera and metabolites are ordered by Ward’s method of hierarchical clustering. Red indicates preferentially ranked for consumption relative to the global community average; blue indicates lower ranked preference.

### BPR scores predict metabolite depletion in held-out bacterial isolates using genus-level models

To evaluate whether genus-level BPR models generalize to new isolates within the same genus, we assessed predictions on eight held-out isolates from *Janthinobacterium, Pseudomonas, Caulobacter, Chryseobacterium, Acinetobacter, Paraburkholderia, Ensifer,* and *Brevundimonas*. These isolates were not included in model training, but other isolates from the same genera were, allowing us to test whether genus-level metabolic representations are sufficient to predict the behavior of novel strains. For each isolate grown in NLDM, we compared the distribution of mean genus-level BPR scores between metabolites that were experimentally decreased (log₂ fold change < −2) and those that were neutral (log₂ fold change ≥ −2) **(Fig 6)**. Across all eight isolates, metabolites with confirmed experimental depletion exhibited significantly higher BPR scores than neutral metabolites (Mann-Whitney U test, p < 0.001 for all genera), demonstrating that genus-level models capture metabolic consumption patterns that transfer to previously unseen strains.

**Figure 6.**
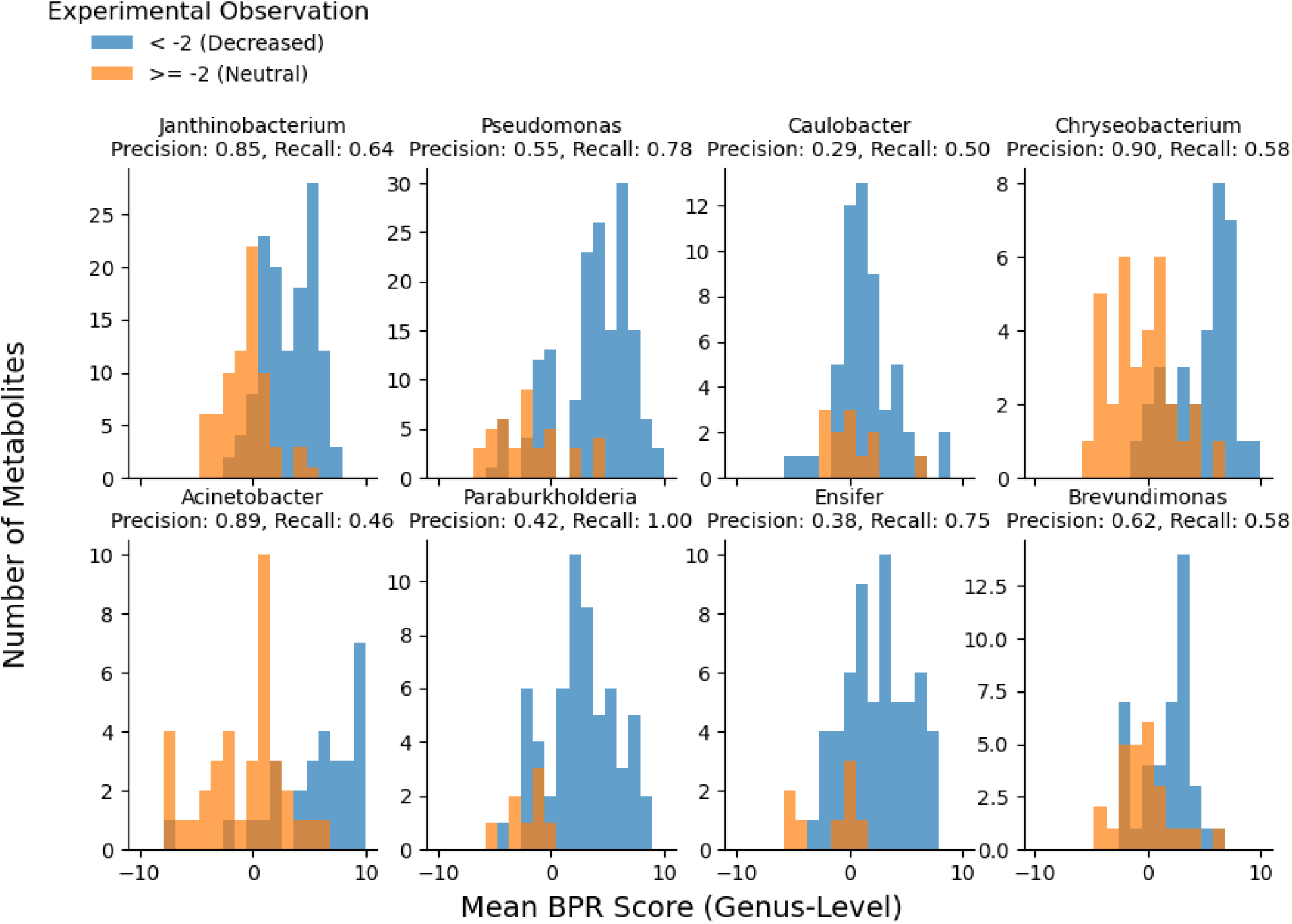
BPR scores discriminate experimentally decreased metabolites from neutral metabolites across diverse bacterial genera for 8 new isolates. Across most genera, metabolites with decreased abundance (log₂ fold change < −2; blue) tend to have higher positive BPR scores, while metabolites with neutral responses (log₂ fold change ≥ −2; orange) shift toward lower or negative scores, indicating that the BPR scoring framework captures biologically meaningful metabolic depletion signatures. Each panel shows overlaid histograms of mean genus-level BPR scores for metabolites measured in NLDM, stratified by experimental outcome. A simple score threshold of zero was used to classify metabolites as predicted decreased (score ≤ 0) or predicted neutral (score > 0), and the resulting precision and recall for each genus are reported above each panel. Precision values are generally high (average ∼0.6), reflecting that metabolites receiving negative BPR scores are predominantly those experimentally confirmed to have neutral responses, though recall varies across genera, indicating differing degrees of sensitivity to true decreases across taxonomic groups.

Using a very simple threshold of zero to classify metabolites as predicted decreased (BPR score > 0) or predicted neutral (BPR score ≤ 0), we observed a median precision of 0.59 and a median recall of 0.61 across the eight isolates **(Fig 6)**. Performance varied considerably: *Chryseobacterium* and *Acinetobacter* achieved the highest precision (0.90 and 0.89, respectively), indicating that positive BPR scores in these genera were highly reliable indicators of true metabolite depletion, though at the expense of moderate recall (0.58 and 0.46). *Janthinobacterium* exhibited a strong balance of high precision (0.85) and moderate recall (0.64). In contrast, *Paraburkholderia* achieved perfect recall (1.00) but lower precision (0.42), suggesting that while the genus-level model captured all truly depleted metabolites, it also assigned positive scores to a substantial fraction of neutral metabolites. *Caulobacter* showed the lowest precision (0.29), indicating that for this genus, the model overpredicted metabolite consumption, possibly reflecting greater metabolic divergence between the held-out isolate and the training isolates. This variation in performance likely reflects differences in intra-genus metabolic diversity: genera with more conserved metabolic profiles across strains are better served by genus-level modeling, while genera harboring greater strain-to-strain metabolic variation will naturally show reduced predictive accuracy.

### Integration of LLM with BPR and growth rate prediction tools to create the WoM Agent

Having found that BPR learned accurate and phylogenetically conserved representations of microbial substrate preferences for most genera, we generalized compound names and taxonomy to enable broader predictions. Compound names were linked to the ChEBI (Chemical Entities of Biological Interest) ontology^22^ using approximate string (“fuzzy”) matching, and microbial isolates were linked to standardized taxonomy, enabling predictions to be aggregated and extrapolated across studies.

We next constructed the full tool using agentic workflow where the LLM formats user inputs, runs the BPR, formats the output, and provides additional context (**Fig. 1**). While we note that an open-weight model exists that has been fine-tuned on soil literature and could serve as the LLM (Qwen2.5-14B^23^), we elected to use a frontier model to avoid the challenges of preventing hallucinations and other unwanted behaviors. We therefore used Gemini 2.5 Flash^24,25^ as the LLM since it is a frontier model that has undergone extensive post-training to help ensure robust behavior.

Substrate preference is only one of many factors that may influence bacterial competition for metabolites (e.g. growth rate, metabolite secretion, and antagonism, among others ^26^). We reasoned that addition of a growth tool would be a valuable complement to BPR since rate determines how fast a microorganism can convert available resources. To do this we used the Phydon model which predicts maximum growth rates directly from genomic sequences ^19^ and operates directly on species or higher level taxonomic phylogeny, making it compatible with our genus level embedding.

### The WoM Agent accurately predicts published time-series substrate depletion

#### Predicting time-series data

We wondered whether the WoM Agent, which uses BPR trained on endpoint exometabolite data, could infer time-series metabolite consumption patterns. To explore this we used a previously published time-series exometabolite profiling study of *Pseudomonas fluorescens* strain FW300-N2E2 grown in a 23-metabolite soil defined medium^28^ noting that the BPR training data included this same strain but grown on a different medium NLDM ^29^ and fructose is not in the BPR model. Model Prompt 1 was used to compare the WoM Agent predictions vs. the LLM alone (Gemini 2.5 Flash).

**Model Prompt 1 (used to generate the results shown in Fig. 7)**

*For Pseudomonas, rank all 23 of these substrates from highest to lowest preference: phenylalanine, glycine, threonine, isoleucine, leucine, lysine, serine, alanine, proline, arginine, glutamate, adenosine, guanine, hypoxanthine, uridine, 4-guanidinobutanoic acid, glucose, fructose, mannose, arabinose, trehalose, maltose, mannitol.*

**Figure 7.**
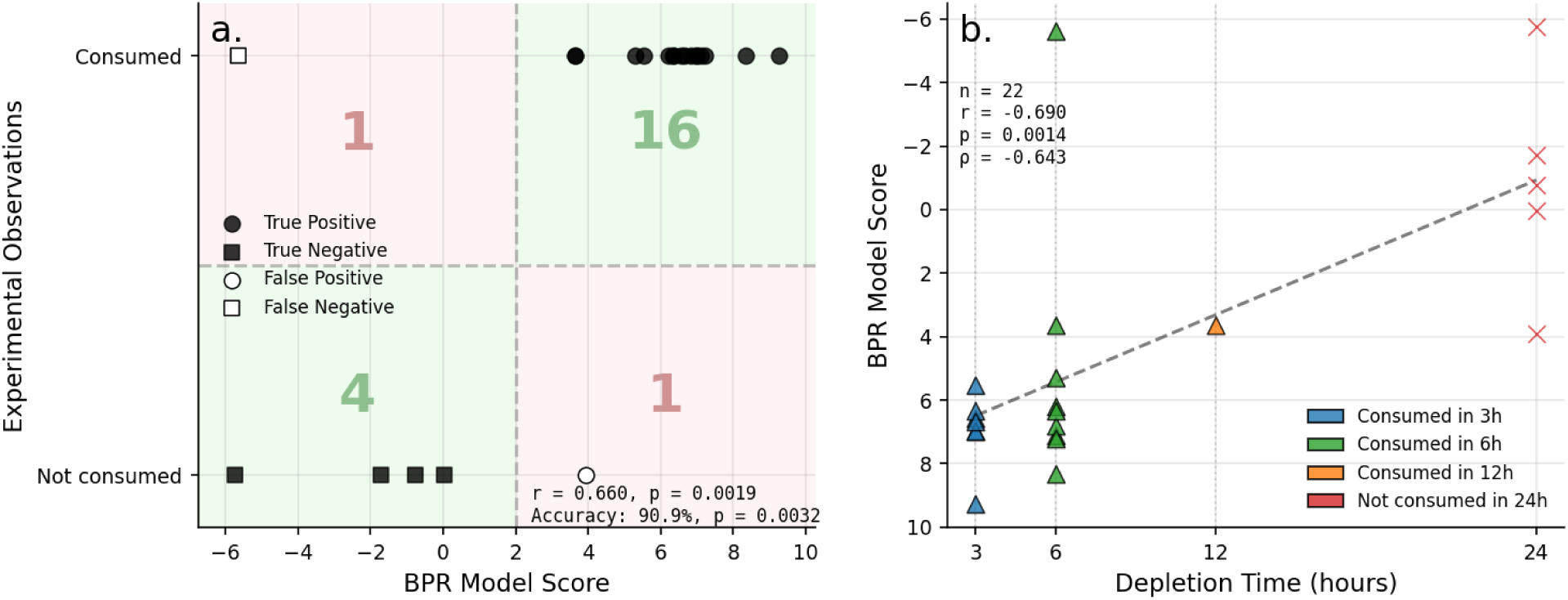
The WoM Agent accurately predicts and interprets substrate consumption and temporal utilization order for *Pseudomonas fluorescens*. a) Predicting metabolite utilization from a growth medium. The BPR model scores for 22 metabolites are compared to experimentally obtained metabolite consumption profile of *Pseudomonas fluorescens* strain FW300-N2E2 derived from exometabolomic data and generated by the Web of Microbes platform^15,28^. b) BPR scores predict substrate preference in time. The BPR model scores for 22 metabolites are plotted against the time (in hours) that more than half of the metabolite was consumed by *Pseudomonas fluorescens* strain FW300-N2E2. Metabolites that were not consumed by more than half in 24 hours were labeled not consumed.

We found that the WoM Agent correctly predicted 16 of the 17 metabolites consumed by the strain (**Fig. 7a**; See SI) based on comparison between the experimental consumption (log2 fold change < −1) and predicted consumption (score > 2). Noting that fructose was absent from the model and no score was generated. Next, we tested whether WoM Agent scores could predict the temporal order of substrate consumption — that is, which metabolites the strain would deplete first from the pool of 22. The time-series study revealed distinct consumption phases, with metabolites depleted during early log phase (3 hours), exponential growth (6 and 9 hours), or not consumed within 24 hours^28^. The scores indeed correlated clearly with consumption timing, with higher scores corresponding to faster consumption (**Fig. 7b**) which is exciting given that the model was not trained-on time-series data. As a control we ran the same prompt on the base LLM Gemini 2.5 Flash. Unlike the WoM Agent, whose predictions are based on deterministic programs (BPR and Phydon), LLM responses are often variable, so we ran the base LLM 10 times. From this we found that it did not predict the correct pattern of usage (See SI: it consistently ranked nucleobases and nucleosides lowest, but the ordering of amino acids relative to sugars varied across runs).

Unlike the base LLM results, the WoM Agent-derived predictions largely matched the experimental data. However, two metabolites did not agree: uridine was predicted to be consumed but was not, and 4-guanidinobutanoic acid was not predicted to be consumed but in fact was. When asked to infer potential reasons for these discrepancies, the agent attributed the 4-guanidinobutanoic acid result to strain diversity and the uridine result to catabolite repression, and both seem reasonable given our experiences (See SI for details).

### The WoM Agent predicts soil bacterial responses to amino acid and xylose addition

#### Soil Spike-in tests

To test the agent’s ability to predict competitive outcomes in a complex environment, we performed soil spike-in experiments, the first using algal amino acids, and the second with xylose. For both experiments soil was amended with one of the two additions or a water control and incubated for 22 and 94 hours. Following incubation, DNA was extracted and characterized using 16S rRNA gene sequencing. The resulting OTU data were aggregated at the genus level and statistical analyses were performed to identify genera that were enriched at either or across time points. The resulting genus-level analyses contained 39 genera present in the WoM dataset.

#### Amino Acid Spike-in model performance

Experimentally we found that two genera, *Pseudomonas* and *Paenibacillus* were enriched (adjusted *p*-values <0.05, SI Table 1). We designed a prompt (Model Prompt 2) asking which of the detected genera (contained in the WoM model) would be predicted to be enriched with the addition of amino acids. The identical prompt was again run on both the WoM Agent and the base LLM. The WoM Agent correctly predicted that *Pseudomonas* would be the most competitive bacterium for the amino acid mixture but did not predict that *Paenibacillus* would also be enriched. Gemini 2.5 Flash was run 10 times and predicted *Pseudomonas* would be the most competitive bacterium 8/10 times and never predicted *Paenibacillus* to be in the top 5 (See SI).

**Model Prompt 2:**

*Make a table where you rank (highest to lowest) of which of these bacteria you predict will be most competitive for a mixture of proteogenic amino acids based on having both a high substrate preferences and high growth rates. Acidovorax, Acinetobacter, Afipia, Arthrobacter, Bacillus, Bradyrhizobium, Brevibacillus, Brevundimonas, Burkholderia, Caulobacter, Chitinophaga, Chryseobacterium, Cupriavidus, Curvibacter, Ensifer, Flavobacterium, Herbaspirillum, Janthinobacterium, Lysinibacillus, Lysobacter, Marmoricola, Massilia, Mesorhizobium, Methylobacterium, Modestobacter, Mucilaginibacter, Niastella, Novosphingobium, Paenibacillus, Pedobacter, Phenylobacterium, Pseudomonas, Rhizobium, Rhodococcus, Sphingomonas, Stenotrophomonas, Streptomyces, Undibacterium, Zoogloea*

**WoM agent response (excerpt, see SI):**

*The Winners: Pseudomonas is predicted to be the most competitive. While Herbaspirillum has a higher raw preference score for amino acids, Pseudomonas grows nearly 6 times faster, allowing it to deplete resources and dominate the population quickly.*

**Gemini 2.5 Flash only response (formatted excerpt, see SI):**

*Predicting bacterial competitiveness relies on a deep understanding of their typical ecological roles, metabolic capabilities, and growth strategies. For proteogenic amino acids, we’re looking for bacteria that are: (1) Metabolically Versatile — can transport and catabolize a wide range of amino acids efficiently; (2) Opportunistic/r-strategists — known for rapid growth when nutrients are abundant; (3) Generalist Heterotrophs — less reliant on highly specialized or recalcitrant substrates. […] Pseudomonas: Top-tier generalist. Extremely metabolically versatile, known for rapid growth rates and efficient degradation of a wide array of organic compounds, including diverse amino acids. Classic opportunistic r-strategist. […] Streptomyces: Excellent degraders, but slower growth. Known for their remarkable ability to break down complex organic polymers, producing amino acids. However, their growth is often slower due to complex life cycles and secondary metabolite production.*

#### Xylose soil spike-in model performance

In addition to the amino acid spike-in study, we performed an additional study where we repeated the same procedure with xylose (Model Prompt 3). Here our experiment found that the only significantly enriched bacterium was from the genus *Novosphingobium* (See SI Table 2). We again prompted both the WoM Agent and base LLM - Gemini 2.5 Flash to determine which of the genera contained in the model would be enriched by the addition of xylose. In this case, only the WoM Agent correctly identified *Novosphingobium*. In the 10 times Gemini 2.5 Flash was run on average *Novosphingobium* was ranked 10th and it was never ranked in the top 5 (See SI).

**Model Prompt 3:**

*Make a table where you rank (highest to lowest) of which of these bacteria you predict will be most competitive for aldehydo-D-xylose based on having both a high substrate preferences and high growth rates. Acidovorax, Acinetobacter, Afipia, Arthrobacter, Bacillus, Bradyrhizobium, Brevibacillus, Brevundimonas, Burkholderia, Caulobacter, Chitinophaga, Chryseobacterium, Cupriavidus, Curvibacter, Ensifer, Flavobacterium, Herbaspirillum, Janthinobacterium, Lysinibacillus, Lysobacter, Marmoricola, Massilia, Mesorhizobium, Methylobacterium, Modestobacter, Mucilaginibacter, Niastella, Novosphingobium, Paenibacillus, Pedobacter, Phenylobacterium, Pseudomonas, Rhizobium, Rhodococcus, Sphingomonas, Stenotrophomonas, Streptomyces, Undibacterium, Zoogloea*

**WoM Agent response (excerpt, see SI):**

*Novosphingobium stands out as a specialist with a score more than double the next closest genus.*

**Gemini 2.5 Flash only response (formatted excerpt, see SI):**

*Pseudomonas: Highly metabolically versatile, often very fast-growing, commonly found in soil and water where plant-derived sugars like xylose are abundant. Many species are known for efficient carbohydrate metabolism. […] Streptomyces: Filamentous Actinobacteria, highly diverse metabolically and known for producing antibiotics. While they can degrade many complex organic compounds, their mycelial growth form often leads to slower overall growth rates compared to unicellular fast growers. […] Bradyrhizobium: Another genus of plant symbionts. Growth rates can be slower than fast-growing saprophytes.*

### The WoM Agent demonstrates autonomous reasoning ability

As a final test of the WoM Agent we examined its autonomous reasoning ability. Specifically, we prompted both the WoM Agent and the base LLM - Gemini 2.5 Flash to propose substrates that could be added to enrich *Streptomyces* vs. *Pseudomonas* given that the latter is an extremely competitive bacterium since it both uses a wide-array of substrates and grows quickly. In contrast, *Streptomyces* are typically slower-growing and more selective. We note that *Streptomyces* is of particular interest given its production of secondary metabolites, making it widely used for biocontrol applications^30^.

**Prompt:**

*I want to selectively enrich Streptomyces in soil without promoting Pseudomonas. Suggest metabolites that Streptomyces strongly prefers but Pseudomonas does not.*

**WoM Agent Top Response (excerpt, see SI):**

*These metabolites represent the best opportunities for selective enrichment:*

*1. Sucrose (Table Sugar)*

*Why: This is your strongest candidate.*

*Streptomyces Score: 3.51 (Strong predicted consumption)*

*Pseudomonas Score: -1.78 (Predicted non-consumption)*

*Note: Many Pseudomonas species lack the invertase enzyme necessary to break down sucrose, whereas Streptomyces are well-known for utilizing diverse sugars.*

*2. Galactitol (Dulcitol)*

*Why: A sugar alcohol derived from galactose.*

*Streptomyces Score: 3.30 (Strong predicted consumption)*

*Pseudomonas Score: -0.77 (Predicted non-consumption)*

*3. Trehalose*

*Why: A disaccharide found in fungi and insects.*

*Streptomyces Score: 2.97 (Good predicted consumption)*

*Pseudomonas Score: 0.04 (Neutral/Weak)*

*Risk: Pseudomonas has a neutral score here, meaning some strains might utilize it slowly, but Streptomyces has a clear metabolic preference.*

**Gemini 2.5 Flash only Top Response (formatted excerpt, see SI):**

*The core principle here is to use complex, recalcitrant polysaccharides that Streptomyces are famous for degrading, and to avoid simple sugars, organic acids, and easily metabolizable compounds that Pseudomonas thrive on.*

*1. **Chitin:** Streptomyces species are renowned for their powerful chitinase enzymes, allowing them to efficiently degrade chitin. While some Pseudomonas species can possess chitinolytic activity, it’s generally not their primary or preferred carbon source, and their efficiency is often significantly lower than Streptomyces. By making chitin the sole carbon source, Streptomyces gets a massive advantage.*

*2. **Xylan (Hemicellulose):** Streptomyces species are well-known for their potent xylanolytic enzyme systems, making them excellent degraders of plant biomass. While some Pseudomonas can utilize components of plant biomass, xylan as a sole complex carbon source will favor the more specialized degraders like Streptomyces.*

The two models used fundamentally different approaches to the problem. The base model responded with general ecological reasoning, characterizing *Pseudomonas* as copiotrophs and *Streptomyces* as oligotrophic polymer decomposers, and recommended complex substrates such as chitin, cellulose, and xylan (none of which are metabolites). While ecologically sound, these recommendations were not grounded in data and focused on polymers outside the scope of small-molecule metabolite amendment specified in the prompt. In contrast, the WoM Agent queried the BPR model and recommended three specific metabolites with quantitative scores: sucrose (*Streptomyces* 3.51, *Pseudomonas* −1.78), galactitol (*Streptomyces* 3.30, *Pseudomonas* −0.77), and trehalose (*Streptomyces* 2.97, *Pseudomonas* 0.04). By contrast, when we run the base LLM 10 times, we find that in runs where sucrose was mentioned (5/10), it explicitly recommended avoiding it, and neither trehalose nor galactitol was suggested in any run. Yet, some of the WoM Agent’s predictions are supported by the literature. For example, wild-type *P. putida* cannot metabolize sucrose natively, requiring engineered expression of invertase and permease genes ^31^, whereas *Streptomyces* species readily utilize sucrose as a carbon source. *Streptomyces* spores accumulate trehalose at up to 25% of dry weight and express inducible trehalose uptake systems^32,33^, while no comparable trehalose catabolism has been reported for *Pseudomonas*. The galactitol prediction represents a less-characterized case where the BPR model generates a testable hypothesis (and even experimental designs, See SI). This comparison illustrates that while the base LLM produces reasonable but variable and generic recommendations, the WoM Agent provides specific, quantitative, and experimentally actionable predictions derived from the underlying BPR model.

## Discussion

In this work we explored whether ranking algorithms widely used for learning human preferences — such as which products or videos a user selects from a bundle of options — could be adapted to learn bacterial substrate preferences from formatted exometabolite profiling data. We compared two factorization methods (SVD and WR-MF) with a pairwise optimization algorithm (BPR) and selected the latter because it uniquely maintained stable leave-one-out performance across all embedding dimensions.

The BPR embeddings captured phylogenetically meaningful trends despite receiving no taxonomic information during training. Substrate preferences were significantly correlated at the genus level, consistent with previous findings on phylogenetic niche conservatism^34,35^. However, we observed numerous departures from strict phylogenetic organization, particularly after removing universally consumed metabolites (**Fig. 5**). These departures may reflect convergent metabolic strategies among distantly related organisms, niche-specific adaptations that cut across taxonomic boundaries, or horizontal acquisition of catabolic capabilities^34^. Important differences at the species level suggest that the model could be substantially improved with additional training data.

The distinctive score analysis revealed metabolites with strong positive scores restricted to one or a few genera, suggesting narrow catabolic specializations, alongside metabolites with broad negative scores across most taxa with a single exception. In some cases, metabolite clustering reflected shared biochemical pathways — for example, structurally related amino acid derivatives grouped together, but often diverse metabolites were clustered together. This structure reflects the degree to which phylogenetically related organisms share distinctive consumption niches in the learned embedding space and is clearly an exciting area for further research.

A validation experiment was performed to test generalization across 8 new isolates (not in the model). The ability of genus-level BPR models to predict metabolite depletion in held-out isolates demonstrates that genus is a meaningful taxonomic unit for modeling bacterial metabolism, with significant separation of BPR score distributions between depleted and neutral metabolites across all eight held-out isolates (*p* < 0.001). This finding has practical implications for the study of microbial communities, where taxonomic assignments are often resolved only to the genus level and strain-level metabolic characterization remains infeasible at scale. At the same time, the variation in precision and recall across genera highlights the inherent limitations of this approach: genera such as *Chryseobacterium* and *Acinetobacter* achieved precision exceeding 0.89, suggesting well-conserved metabolic repertoires, while lower precision for *Caulobacter*, *Paraburkholderia*, and *Ensifer* likely reflects greater intra-genus metabolic diversity, where the model overpredicts consumption by attributing the metabolic capabilities of training isolates to the held-out strain. For applications requiring higher accuracy, or for genera with substantial strain-to-strain metabolic heterogeneity, incorporating sub-genus information such as species-level assignments, phylogenetic placement, or genomic content could improve predictions, but the framework presented here establishes a strong baseline demonstrating that genus-level metabolic modeling provides actionable predictions even in the absence of strain-specific training data.

Integration of BPR with an LLM made the model accessible through natural language prompts and enabled the agent to provide biological context alongside predictions. Validation against published data demonstrated the agent’s ability to generalize beyond its training conditions. BPR accurately predicted substrate consumption by *Pseudomonas fluorescens* FW300-N2E2 in a medium not used for training, correctly identifying 16 of 17 consumed metabolites, indicating generalization across media. More strikingly, BPR scores correlated with the temporal order of consumption, despite being trained only on endpoint data — suggesting that the learned preference rankings reflect an underlying hierarchy of substrate utilization that manifests as sequential depletion. The WoM Agent also uniquely predicted the rapid consumption of hypoxanthine and nucleotides, which the base Gemini model expected to be among the last substrates used, highlighting the critical contribution of the BPR model to the agentic workflow.

A soil spike-in experiment was used as an initial test of the agent’s ability to predict competitive outcomes in a complex natural environment. The agent correctly predicted that *Pseudomonas* would be the most competitive genus (of those in the model) following amino acid addition, and *Novosphingobium* would respond to xylose. The amino acid result is what would be expected both from the literature and was also predicted by the base LLM. When all 600 OTUs detected were considered, a second strain of the *Tumebacillus* genus was also found to be significantly increased, though this was not considered by the agent since it isn’t represented in the model. However, in the case of the xylose, the WoM Agent was uniquely able to predict that *Novosphingobium* would be the most competitive and this was the only isolate found to be enriched across all OTUs. We note that our spike-in study was performed using low levels of substrate and incubations were short lived inside small tubes, all which can impact the results.

We also compared the base LLM and the WoM Agent on autonomous reasoning for the highly relevant use case of selectively promoting the growth of *Streptomyces* over *Pseudomonads*. *Streptomyces* are thought to be beneficial biocontrol organisms and so molecules that can promote their growth could have important applications in plant health^36^. However, they must compete with organisms such as *Pseudomonads* that have both high growth rates and broad substrate uses. We found that the base model drew on broad training knowledge to produce ecologically reasonable but generic recommendations, essentially recapitulating textbook understanding of *Streptomyces* and *Pseudomonas* ecology to recommend the biopolymers chitin, starch, and inulin–however, none of which are metabolites as specified in the prompt. In contrast, the WoM Agent generated specific predictions for metabolites: two of which had literature support (sucrose and trehalose) and a novel testable hypothesis for galactitol. We see the LLM’s polymer recommendations and the agent’s small-molecule predictions as complementary, suggesting that future iterations of the agent could benefit from also allowing the LLM to make independent predictions that can subsequently be integrated in the final Agents response. Thus taking advantage of the ongoing improvements in frontier LLMs while also grounding it in determinant models.

We acknowledge several important limitations. While outcomes in natural environments depend on metabolite secretion, antagonism, spatial structure, and many other factors, the current WoM Agent only has tools for substrate preference and growth rate predictions. While we find that for such a simple model, BPR performs surprisingly well, it has limited accuracy and likely generalization across possible use cases and so results should be interpreted with caution. BPR is primarily trained on defined media and while this has the advantage that exometabolomics is able to monitor ‘all choices’ it will be important to extend this to more complex media that may include undetected metabolites. Likewise, given the importance of cross-feeding and secondary metabolites in microbial ecology, it will be important to add tools for modeling metabolite secretion. The BPR model’s accuracy varies taxonomically and we have already identified some problematic clades. However, accuracy will also depend on representation in the training data and so predictions for taxa or compounds far from the training distribution should be interpreted with caution. Furthermore, the current model treats substrate preference as a fixed trait, whereas in reality they may vary with environmental conditions, community context, and growth phase. We also recognize that the Phydon model used for growth rate prediction assumes translation is the bottleneck. While this may often be true especially for copiotrophic microbes growing under replete conditions, it likely is not under low-growth conditions. Finally, we note that it is challenging to engineer the LLM system prompt to ensure that it consistently uses the BPR and Phydon tools vs. its own tendency to provide an immediate response so it is advisable to confirm that the output is based on the tools. As is typical with LLMs we’ve observed that identical prompts can result in slightly different contextual responses.

Despite these limitations, the WoM Agent demonstrates that ranking algorithms borrowed from recommender systems can capture meaningful patterns in microbial substrate preferences and that integration of BPR and Phydon models as tools for an agentic LLM workflow makes them broadly useful and accessible. The addition of more isolates, metabolites, and metabolite secretion should greatly improve performance. Likewise adding more tools such as those based on genomic analysis, mutant fitness, etc will make the agent more accurate and robust.

## Conclusion

The WoM Agent demonstrates that integrating preference and growth models with LLM-driven orchestration creates an accessible, extensible tool for exploring microbial substrate use. We found that a Bayesian Personalized Ranking (BPR) approach learned accurate and taxonomically conserved patterns of substrate preference from exometabolite profiling data which is to our knowledge, the first such application of preference ranking algorithms. The WoM Agent uses a LLM to interpret natural language prompts, execute both BPR and Phydon growth models, and help users interpret results in biological context. The agent accurately predicted published time-series substrate use by *P. fluorescens* in a medium not used for training, where the base LLM alone could not, and correctly predicted soil bacteria stimulated by metabolite additions. We recognize that the current agent does not account for genomics, complex ecosystem processes and so predictions should be interpreted with caution. Important future directions include greatly expanding the exometabolite training data and providing the agent with additional tools to improve prediction accuracy across complex use cases, from predicting prebiotics for microbiome engineering to generating new hypotheses in microbial ecology.

## Methods

### Exometabolite Dataset Construction

Binary metabolite consumption and secretion profiles were derived from exometabolomic data generated by the Web of Microbes platform^15^ (Northen Lab, Lawrence Berkeley National Laboratory, Feb 2026). The raw data consist of organism-by-metabolite matrices in which each cell contains a numerical score reflecting the change in abundance of a given metabolite in spent medium relative to uninoculated controls, with negative values indicating net consumption and positive values indicating net release. Where time-series measurements were available for the same isolate, only the final time point was retained to capture endpoint consumption and secretion behavior.

Compound names from the exometabolomic source files were standardized against the ChEBI (Chemical Entities of Biological Interest) database ^22^. ChEBI compound records, synonyms, and structural data were downloaded and processed to build a reference name dictionary. Each compound name in the exometabolomic data was normalized by converting to lowercase and stripping whitespace, hyphens, underscores, and mass spectrometry adduct annotations (e.g., [M+H]+). Several known ambiguous compound names — including stereoisomer mixtures and co-eluting species — were manually mapped to a single representative entry prior to automated matching. Normalized names were then matched to ChEBI entries first by exact string comparison and, if no exact match was found, by fuzzy string matching using the RapidFuzz library (token ratio scorer, minimum score threshold of 90). Compounds were further deduplicated by the first block of their InChI key, which encodes molecular connectivity independent of stereochemistry, so that stereoisomers of the same compound were collapsed to a single representative entry. Compounds that could not be matched to any ChEBI entry at the 90% similarity threshold were excluded from the dataset.

Numerical exometabolomic scores were converted to categorical assertions: metabolites with scores below −2 were classified as consumed, scores above +2 as secreted, and intermediate values as neutral. Metabolites consumed by fewer than three isolates across the entire dataset were removed to exclude rarely observed or potentially spurious interactions.

Taxonomic assignments for each microbial isolate were obtained from a curated phylogeny table providing genus, family, order, class, and phylum ranks. This taxonomy was joined to the exometabolomic data so that each observation is annotated with both the metabolite identity and the full taxonomic lineage of the organism.

The final dataset consists of organism–metabolite pairs annotated with binary consumption and secretion labels, standardized compound identities linked to ChEBI, and taxonomic classifications at all major ranks. This dataset serves as input to all subsequent modeling steps.

### Matrix Factorization Models

We framed the prediction of microbial metabolite consumption as an implicit feedback recommendation problem, in which each organism’s set of consumed metabolites constitutes its positive interactions and all other metabolites are unobserved rather than explicitly negative. The binary organism-by-metabolite interaction matrix was factorized into low-dimensional embedding vectors for each organism and each metabolite, such that the dot product between an organism’s embedding and a metabolite’s embedding approximates the likelihood of consumption. We compared three factorization approaches, as described below.

Bayesian Personalized Ranking (BPR) optimizes a pairwise ranking objective: for each organism, observed positive metabolites should receive higher scores than unobserved metabolites ^17^. Organism and metabolite embeddings are represented as PyTorch embedding layers initialized with Gaussian noise (standard deviation 0.01). Training proceeds by stochastic gradient descent using the Adam optimizer (learning rate 0.01), sampling 256 triplets per batch for 1000 epochs. Each triplet consists of an organism, a randomly selected positive metabolite, and a randomly sampled negative metabolite (sampled uniformly from all metabolites not in the organism’s positive set). The loss for each triplet is the negative log-sigmoid of the score difference between the positive and negative items, averaged over the batch.

Weighted Regularized Matrix Factorization (WR-MF) treats the binary interaction matrix as implicit confidence data and solves for embeddings using alternating least squares ^21^. We used the implicit library implementation with L2 regularization strength 0.1 and 50 ALS iterations. The sparse organism-by-metabolite matrix was constructed with confidence weights of 1.0 for all observed positive interactions.

Truncated Singular Value Decomposition (SVD) serves as a non-iterative baseline. The binary interaction matrix is factorized directly using scikit-learn’s TruncatedSVD ^20^ implementation, yielding left singular vectors as organism embeddings and right singular vectors as metabolite embeddings. No weighting or regularization is applied.

All three methods produce organism embeddings and metabolite embeddings of matched dimensionality, and predictions for any organism-metabolite pair are computed as the dot product of the corresponding vectors. We evaluated embedding dimensions of 4, 8, 16, 32, 64, and 128. Based on the leave-one-out cross-validation results (Fig 1 a,b), we selected BPR with 32-dimensional embeddings for all subsequent analyses.

### Genus-Level Distinctive Preference Heatmap

To identify metabolites whose consumption distinguishes specific taxa from the broader microbial community, we constructed a genus-by-metabolite heatmap of distinctive scores. Genera were ordered along the horizontal axis according to their taxonomic hierarchy, sorted alphabetically within class, order, family, and genus ranks (color bar indicating class membership). A genus embedding was computed as the mean of all member isolate embeddings, and distinctive scores were calculated for each genus-metabolite pair as described above. Metabolites classified as universally consumed (global prevalence exceeding 0.7 across all isolates) were excluded from the visualization to focus on compounds that differentiate taxa. Among the remaining metabolites, the 60 with the highest variance in distinctive score across genera were retained, selecting for compounds with the greatest discriminatory power between taxonomic groups. Metabolites and genera were ordered by Ward’s method. The heatmap uses a diverging red-blue color scale centered at zero, where red indicates metabolites that a genus preferentially consumes relative to the global community average and blue indicates relative avoidance.

To quantify the degree to which substrate preferences are conserved at the genus level, we compared the cosine similarity of embedding vectors for isolate pairs within the same genus to pairs from different genera using a one-sided Mann-Whitney U test. We additionally performed a permutational multivariate analysis of variance (PERMANOVA) on the Euclidean distance matrix of isolate embeddings with genus as the grouping factor (999 permutations) to estimate the fraction of total embedding variance attributable to genus identity. As a complementary measure operating on predicted preferences rather than raw embeddings, we computed pairwise Spearman rank correlations of the full metabolite score vectors (isolate embedding dotted with all metabolite embeddings) for within-genus and between-genus pairs and compared their distributions with a Mann-Whitney U test. All analyses were restricted to genera containing at least two isolates.

#### Distinctive Score Calculation

The raw preference scores produced by the BPR model reflect both taxon-specific metabolite preferences and a global popularity signal shared across all organisms — metabolites that are widely consumed by most microbes naturally receive high scores regardless of the queried taxon. To separate these two components, we computed a distinctive score that isolates the taxon-specific signal by removing the contribution of global popularity from each taxon’s predicted preferences.

We first computed a global direction vector by averaging all microbe embedding vectors in the trained BPR model and normalizing the result to unit length. This vector represents the “average organism” in embedding space, and its dot product with each metabolite embedding yields a global popularity score reflecting how universally preferred each metabolite is across the full community. To obtain the distinctive score for a given taxon, we constructed a taxon embedding by averaging the learned embeddings of all isolates belonging to that taxon, then projected out the global direction by subtracting the component of the taxon embedding that lies along the global direction vector (i.e., computing the residual via v_distinctive = v_taxon − (v_taxon · ĝ)ĝ, where ĝ is the unit global direction). The distinctive score for each metabolite is then the dot product of the metabolite embedding with this residual taxon vector. Positive distinctive scores indicate metabolites that the taxon preferentially consumes relative to the community average, while negative scores indicate metabolites the taxon avoids relative to expectations from global popularity alone.

### Taxonomic Holdout Cross-Validation

To evaluate whether the model’s learned embeddings generalize across phylogenetic distances, we performed a hierarchical holdout benchmark in which entire taxonomic groups were withheld from training and predictions for held-out isolates were derived from the embeddings of their nearest taxonomic relatives remaining in the training set. We evaluated four holdout levels: order, family, genus, and isolate. Thus, representing increasing phylogenetic proximity between training and test organisms. At each level, every group at that taxonomic rank was held out in turn: all isolates belonging to the focal taxon were removed from the training data, a new BPR model (embedding dimension 32) was retrained on the remaining organisms, and predictions were generated for each held-out isolate. The prediction embedding for each held-out group was computed as the mean embedding of non-held-out organisms belonging to the same taxon one rank higher in the hierarchy. Specifically, when holding out a genus, the prediction was derived from other genera in the same family; when holding out a family, from other families in the same order; when holding out an order, from other orders in the same class; and when holding out an individual isolate, from other isolates in the same genus. Groups for which no relatives remained in the training set at the next taxonomic level were excluded from evaluation.

For each held-out isolate, predicted metabolite scores were computed by taking the dot product of the prediction embedding with all metabolite embeddings. Two scoring variants were evaluated: raw scores using the prediction embedding directly, and distinctive scores using the prediction embedding after projecting out the global direction (the unit vector representing the mean of all training microbe embeddings), as described above. Performance was assessed using two metrics. The mean percentile rank measures the average position of each isolate’s true positive metabolites within the full ranked prediction list, where lower values indicate better performance and 0.5 represents random expectation. The AUC was computed as the fraction of negative metabolites ranked below each true positive, averaged across all positives for each isolate, where 0.5 again represents chance performance. Results are presented as distributions across all held-out isolates at each holdout level, with raw and distinctive scores compared side by side to assess whether removing the global popularity signal improves or degrades generalization at different phylogenetic distances.

### BPR validation experiment

### Bacterial culturing

Northen Lab Defined Medium^29^ (NLDM) was used for all culturing experiments following a published protocol^27^. We selected 8 new isolates that were not contained in the model that spanned different genera: *Acinetobacter* sp., *Brevundimonas* sp., *Caulobacter* sp., *Chryseobacterium* sp., *Ensifer* sp., *Janthinobacterium* sp., *Paraburkholderia* sp., *Pseudomonas* sp. For exometabolomics measurements, precultures were first grown in NLDM, R2A or LB at 30°C (150-200 rpm). Once sufficient biomass was obtained, preculture OD_600_ values were taken so that calculations could be made such that the subsequent 2.0 mL exometabolomic culture could be inoculated to a starting OD_600_ of 0.01-0.02. An appropriate volume was removed from the preculture and transferred to a sterile 1.5 mL microcentrifuge tube and centrifuged at 7,500 xg for 5 minutes at 4°C. Following, the media was decanted and the pellet was resuspended in 1 mL of sterile 1x PBS to wash the cell pellet. After, the cells were pelleted again by centrifugation at 7,500 xg for 5 minutes at 4°C. The PBS washing was conducted twice more. Following, the cell pellet was resuspended into an appropriate volume of culture medium to inoculate the exometabolomic culture to 0.01-0.02 OD_600_. Cultures were run as triplicates on 24 well plates with 2 mL culture volumes, using an INFORS HT multitron pro set to 650 rpm and 30°C. Each plate was composed of uninoculated NLDM (3 total samples) that were used as the control reference for fold-changes per metabolite per experiment. After 24 hours of growth, OD_600_ was measured for the plate and 1000 uL of each well was taken and centrifuged at 7,500 xg for 5 minutes at 4°C. After centrifugation, 850 μL of the media was transferred to a fresh sterile microcentrifuge tube and stored at -80 C until preparation for LC-MS.

### Metabolite analysis

The supernatant samples were frozen at −80°C and then freeze-dried (Labconco Freeze-Zone). The dried material was resuspended in 150 μl of LC-MS grade methanol containing internal standards^40^. The solution was vortexed 10 seconds twice, bath-sonicated in ice water for 15 minutes, and centrifuged (10,000 xg for 5 minutes at 4°C) to pellet insoluble material, and then supernatants were filtered using 0.22-μm polyvinylidene difluoride microcentrifuge filtration devices (Pall) (10,000 xg for 5 minutes at 4°C). Metabolites were separated using hydrophilic interaction liquid chromatography (HILIC) for polar metabolomics. Analyses were performed using an InfinityLab Poroshell 120 HILIC-Z column (Agilent) on an Agilent 1290 HPLC connected to a Q Exactive hybrid quadrupole-Orbitrap mass spectrometer (Thermo Fisher Scientific) using ElectroSpray Ionization (ESI). LC-MS/MS and ESI parameters are previously published^40^.

### Evaluation of genus-level BPR model predictions on held-out isolates

To assess whether genus-level BPR models generalize to new isolates, we evaluated predictions on eight bacterial isolates excluded from model training (*Janthinobacterium, Pseudomonas, Caulobacter, Chryseobacterium, Acinetobacter, Paraburkholderia, Ensifer, and Brevundimonas*). The exometabolomic data described above was used to classify metabolites as experimentally decreased (log₂ fold change < −2) or neutral (log₂ fold change ≥ −2). Mean genus-level BPR scores were compared between decreased and neutral metabolites using the Mann-Whitney U test, and a binary prediction was generated by thresholding at zero, with scores greater than zero predicted as decreased. Precision and recall were computed using scikit-learn, and all analyses were performed in Python using NumPy, SciPy, and scikit-learn.

### Soil spike in experiment

Soils were collected from Plots 5 and 12 from the Angelo Range Reserve using a 2” diameter corer in 10 cm increments from 0 – 20 cm^37^. The samples at each depth and sampling location were transferred to sterile whirlpacks and kept on ice until transported into the lab where they were stored at 4°C. Each soil was sieved (2 mm) and stored at 4°C in sterile 50 ml conical tubes until experimented on. 250 mg of soils were aliquoted into 2 ml microcentrifuge tubes and incubated overnight at 18 °C to equilibrate and then spiked with 60 ul of 200 μM of ^13^C-xylose (Omicron Biochemicals, N=3) or ^13^C-labeled algal amino acids (Cambridge Isotopes, N=3) which were supplied as so that the most abundant compound in the mix, 13C alanine was at 200 μM. All compounds were supplied in a ‘synthetic rainwater’ which was formulated based on data collected by the National Atmospheric Deposition Program for their Hopland, CA station: contained the following trace ions (in μM at final concentration): 0.935 Mg^2+^, 0.86 Ca^2+^, 0.38 K^+^, 6.9 Na^+^, 4.41 NH ^+^, 4.15 NO ^-^, 8.12 Cl^-^, 1.7 SO ^2-^. Controls were synthetic rainwater only. For community composition analysis, samples were removed at 22 and 94 hrs and the contents of a Mo Bio Power Soil disruption tube were added to each sample tube, it was then vortexed and frozen at -80°C until extraction following manufacturer’s instructions except that bead beating was performed using a Qiagen TissueLyser II at 30 HZ for 10 minutes. Extracted DNA was normalized and then V4 regions were sequenced as described previously^38^. 16S OTU tables were prepared as described earlier and taxonomic assignment was performed in Usearch using the SILVA v123 LTP named isolate subset (https://www.drive5.com/usearch/manual/sintax_downloads.html)^38^. Genus and family-level sample tables were produced by clustering by taxonomic assignment and statistical analysis was performed using the DESeq2 Software^39^.

## Data Availability

All WoM formatted data and scripts are provided here: https://github.com/northenlab/webofmicrobes

## Code Availability

https://github.com/northenlab/webofmicrobes

## Acknowledgements

We thank Valentine Trotter for providing some of the strains used in this study. M.W. gratefully acknowledges start-up support from UC Riverside for his contributions. The other authors acknowledge support from Ecosystems and Networks Integrated with Genes and Molecular Assemblies (ENIGMA, http://enigma.lbl.gov) and Microbial Community Analysis & Functional Evaluation in Soils (m-CAFEs, https://mcafes.lbl.gov) Science Focus Areas led by Lawrence Berkeley National Laboratory as well as the Enhancing Camelina Oilseed Production with Minimum Nitrogen Fertilization in Sustainable Cropping Systems (ECON), and this work also used resources of the National Energy Research Scientific Computing Center (NERSC, https://www.nersc.gov/)--all of which are supported by the U.S. Department of Energy, Office of Science, Office of Biological & Environmental Research under contract number DE-AC02-05CH11231 to Lawrence Berkeley National Laboratory.

## AI Declaration

Claude Opus 4.5 was used for refining text. Gemini 2.5 Flash was used for experiments as described in the methods section.

## Author Contributions

Benjamin P. Bowen - B.P.B. Conceptualization, final model, model integration, original draft, reviewing and editing, and figures.

Markus de Raad- M.d.R. Data generation, analysis figures, and writing.

Suzanne M. Kosina - S.M.K. Data generation, analysis, and reviewing and editing.

Peter F. Andeer - P.F.A. Performed the soil experiments and reviewing and editing.

Vlastimil Novak - V.N. reviewing and editing.

Katherine B. Louie - K.B.L. Data generation and writing.

Bradley Biggs- B.B. - Data generation and reviewing and editing.

Timothy Paulitz–T.P. - Data generation.

Hao Peng–H.P. - Data generation.

Mingxun Wang - M.W. Performed the LLM integration and UI.

Adam Arkin- A.P.A. Data generation and reviewing and editing.

Trent R. Northen - T.R.N. Conceptualization, initial prototyping, figures, original draft, reviewing and editing.

